# New annotations for three pea aphid genome assemblies allow comparative analyses of duplication and gene family evolution

**DOI:** 10.1101/2025.05.08.652899

**Authors:** Kevin D. Deem, Jennifer A. Brisson

## Abstract

Reliable genome annotation is crucial for analyses of gene function, conservation, duplication, and evolution. Factors such as the sequencing technology used to create the assembly, as well as duplication and rearrangements within the genome of interest, can have a large impact on the quality of gene annotations. In particular, short-read-based assemblies tend to mis-assemble duplicated genes as single loci, a problem that requires additional long-read sequencing to resolve. Pea aphids exhibit a high level of gene duplication from frequent genomic rearrangements, which has led to the mis-assembly and mis-annotation of genes. Here, we re-annotate the pea aphid reference genome, along with two long-read pea aphid genomes, to facilitate future analyses of gene duplication and function in pea aphids. We use an integrated approach, consolidating both *ab initio* and RNA-Seq-based annotations into unified gene models. The new annotations contain genes that were missing, mis-annotated, or mis-assembled in the reference genome, and are generally consistent across assemblies, showing very good agreement between the long-read assemblies. Our annotation method is sensitive enough to refine existing gene models, uncovering alternatively used promoters and isoforms, and aids in finding gene duplications. These data provide a useful supplement to the existing reference annotations and a new comparative framework for discovery and analysis of gene function and duplication in this important emerging model insect.

## Introduction

Proper assembly and annotation of genomes is crucial for bioinformatic and molecular biology investigations into gene function and evolution. As arguably one of the most important processes in gene family evolution, gene duplication presents a unique challenge for the proper assembly and annotation of genomes. The method(s) by which a genome assembly is produced have a substantial impact on the accuracy of gene duplicate assembly and annotation. One major deciding factor in accuracy of assembly is the choice of read type. Illumina short reads (50-300bp) are readily obtained in high numbers in a cost-effective manner. However, short-read-based genome assemblies are the most prone to mis-assemble duplicated genes, especially for recent duplications (Alkan et al., 2011; Mastretta-Yanes et al., 2014; Wang et al., 2021). Long-read technologies such as Pacific Bioscience (Eid et al., 2009) or Oxford Nanopore Technologies (Jain et al., 2016), provide the most accurate assembly of overall gene structure, even in the trickiest cases of recent or partial gene duplication (Fadaie et al., 2021; Logsdon et al., 2020; Nguyen, 2024; Showpnil et al., 2024).

Pea aphids (*Acyrthosiphon pisum*) are an important insect model for studying gene duplication (Fernández et al., 2020), the molecular genetic basis of morphological evolution (Li et al., 2020), and aphid-bacterial symbiont interactions (Price et al., 2011). However, these types of studies can be hindered by mis-assembly within the short-read pea aphid reference genome (Li et al., 2019). For instance, new long-read genome assemblies from winged and wingless males were required to properly assemble a region on the X chromosome controlling male wing dimorphism called *api* (Braendle et al., 2005; Li et al., 2020). This region was mis-assembled in the short-read reference genome (Li et al., 2019), but the assembled long-read genome revealed the presence of a duplicated gene (*follistatin-3*) as part of an insertion specific to wingless males (Li et al., 2020). More recently, we investigated aphid-specific duplications of wing gene regulatory network components and found that a number of these newly identified paralogs were incorrectly annotated in the reference genome (Saleh Ziabari et al., 2025a). These instances of mis-assembly and mis-annotation highlight the need for higher quality genome assemblies and annotations in this species.

Multiple methods exist for gene annotation, each with their own advantages and drawbacks. *Ab initio* methods such as AUGUSTUS (Stanke and Morgenstern, 2005) allow annotation of genes from raw genome sequence by predicting groups of exons that code for proteins. While these methods require no additional data beyond the assembled genome to perform gene annotation, the accuracy and completeness of the annotation may suffer as a result. This accuracy can be increased by providing RNA-Seq data to better train models of splice sites using BRAKER1 (Hoff et al., 2016). Other RNA-Seq based methods of transcript discovery such as genome-guided Trinity (Grabherr et al., 2011) or StringTie (Pertea et al., 2015) utilize read sequence and/or coverage to identify exons and intron-spanning reads to determine how they are spliced into transcripts. These methods can be highly accurate and provide more complete gene models, but they can only identify genes expressed in RNA-Seq input (*i*.*e*. tissue-specific data will provide fewer genes than whole-body data). In addition, Trinity can also assemble transcriptomes *de novo* purely from RNA-Seq reads (the initial intended use of the software), potentially uncovering genes that are missing or mis-assembled in a given genome (Grabherr et al., 2011).

The software PASA takes a more unified approach, integrating Trinity *de novo*, Trinity genome-guided, BRAKER1, and StringTie transcript predictions into clustered gene models (Haas et al., 2008). These transcripts are first filtered for protein coding potential, quality of alignment to the genome, and intron size before being clustered into genes based on overlap (Haas et al., 2008). The advantage conferred by using PASA is that the complementary strengths of *ab initio* and RNA-Seq-based methods work synergistically when combined into a unified annotation.

Here we worked to develop a comprehensive reference annotation for the pea aphid genome. To do this, we integrated RNA-Seq data from many morphs (asexual females, sexual females, males; winged and wingless when available) and developmental stages (embryos, nymphal instars, and adults) with the reference genome (Li et al., 2019), as well as our own long-read assemblies (Li et al., 2020). We then used Trinity (genome-guided and *de novo*), StringTie, and BRAKER1 to individually predict transcripts, followed by PASA clustering of transcripts into unified gene models. We have made our long-read assemblies, a version of the reference assembly with a corrected *api* region, and their new annotations publicly available. These new gene annotations provide a rich source of information for future studies of duplication, function, and gene family evolution in pea aphids and other insects.

## Methods

A summary diagram of the workflow described below is provided (**Supplemental Figure S1**).

### Generation of winged and wingless male embryo transcriptomes

Pea aphid mothers producing winged and wingless males were generated as previously described (Saleh Ziabari et al., 2025b). Briefly, asexual females from lines producing exclusively winged (412) or wingless (409) males were reared in short day conditions to induce production of males and sexual females. Once mothers stop producing females, they begin producing only males, which was confirmed via PCR as in Saleh Ziabari et al. (2025b). Embryos were collected from the ovaries of male-producing mothers and sorted into young (germaria through onset of eye pigmentation) and old (onset of eye-pigmentation through birth) samples. Total RNA was extracted from embryos via TRIzol-chloroform extraction (Invitrogen, Carlsbad, CA) and treated with DNaseI. RNA clean-up was performed on 3ug of total RNA per replicate with the Zymo Clean & Concentrator-5 kit (Zymo Research, Irvine, CA). Three biological replicates of each of the four conditions (young and old wingless, young and old winged) were then sequenced on an Illumina HiSeq 3000 producing 20-30M 2×150bp paired-end reads per replicate (Azenta Life Sciences, Burlington, MA). Raw reads were submitted to NCBI under BioProject PRJNA1380473.

### Trinity *de novo* transcriptome assembly

Only paired-end reads were used to generate the Trinity *de novo* assembly (Grabherr et al., 2011). Reads were first adapter- and quality-trimmed with Trimmomatic v0.39 (Bolger et al., 2014) in paired-end mode using the parameters “ILLUMINACLIP: TruSeq3-PE-2.fa:2:30:10 SLIDINGWINDOW:4:5 LEADING:5 TRAILING:5 MINLEN:25”. Reads were then randomly subsampled to 2% using the view method in Sambamba v0.6.6 (Tarasov et al., 2015) with the parameter “-s 0.02”. Trinity (v2.15.1) *de novo* assembly of transcripts was then performed in singularity-ce v3.11.4 (Kurtzer et al., 2017) using the image file trinityrnaseq.v2.15.1.simg and the parameters “--normalize_by_read_set --no_parallel_norm_stats --min_kmer_cov 2”.

### Read mapping and RNA-Seq-based transcript assembly

Both single-end and paired-end samples were used for RNA-Seq-based gene annotation methods. Read accessions (Grantham et al., 2020; Purandare et al., 2014; Saleh Ziabari et al., 2025b) are found in **Supplemental Table S1**. All of the following steps were performed on the Winged Male Long-Read, Wingless Male Long-Read (Li et al., 2020), and what we call the Modified Reference (Li et al., 2019; Saleh Ziabari et al., 2025b) genome assemblies. For the latter, we modified the reference genome of (Li et al., 2019) to include an insertion that is present solely in wingless males (Li et al., 2020). Specifically, we soft-masekd positions 62,565,000-62,660,000 on the X chromosome of that genome (Li et al., 2019) and added a new scaffold containing the properly assembled *api* insertion region from the Wingless Long-Read assembly (Li et al., 2020) named “NC_WLinsert_pilon10_start62575000_to_62660000” (Saleh Ziabari et al., 2025b).

Reads were aligned to each assembly with Hisat2 v2.2.1 (Kim et al., 2019) using default mapping parameters. The .sam files for each replicate from each sample were converted to .bam, sorted, merged, and indexed using SAMtools v1.13 (Li et al., 2009). Duplicate reads were removed from the merged replicate .bam files using the markdup function of Sambamba v0.6.6 with the parameter “--remove-duplicates”. These files were then each randomly subsampled at a rate of 20% using the view method of Sambamba with the parameter “-s 0.2”. These merged-replicate, duplicate-less, subsampled .bam files were then merged using SAMtools and used for all RNA-Seq-based annotation methods. Braker was run in ET-mode (Hoff et al., 2016) on the respective processed .bam file for each assembly in singularity-ce v3.11.4 (Kurtzer et al., 2017) using the image braker3.sif (Gabriel et al., 2024). Genome-guided transcriptome assembly was performed with these files for each assembly in Trinity v2.15.1 (Grabherr et al., 2011), also run in singularity-ce v3.11.4 with the image trinityrnaseq.v2.15.1.simg, using the flag “--genome_guided_bam” and the parameter “--genome_guided_max_intron 200000”. Transcripts were mapped to the genome using GMAP v2021-08-25 (Wu and Watanabe, 2005). StringTie v2.2.3 (Pertea et al., 2015) was run using default parameters and transcript sequences were extracted from genomic intervals using GffRead v0.12.7-2 (Pertea and Pertea, 2020).

### Integrated gene clustering of transcripts using PASA

Transcript files from Trinity *de novo* (Trinity-DN) and genome-guided (Trinity-GG) assembly, and genomic coordinate files output from StringTie and Braker, were filtered and clustered into genes using PASA v2.5.3 (Haas et al., 2008) for all three assemblies. Following extraction of Trinity-DN accessions, clustering was run in singularity-ce v3.11.4 using the image file pasapipeline.v2.5.3.simg with the following parameters: “--create --run -I 100000 --ALIGNERS gmap --ALT_SPLICE –TRANSDECODER”. The final output included nucleotide sequences and genomic intervals for high-confidence protein-coding transcript isoforms with a maximum intron size of 100kb clustered into ‘genes’.

### Protein prediction, BUSCO analysis, and ortholog clustering

Protein sequences coded by transcripts from Trinity-DN, and from Trinity-GG, StringTie -> GffRead, Braker, and PASA for each genome assembly, were predicted separately using the “Longest-ORFs.fasta” output of TransDecoder (Haas, n.d.) run either locally or on the Galaxy webserver (The Galaxy Community et al., 2024). Gene-transcript maps output from Trinity-DN and Trinity-GG were provided using the “–gene_trans_map parameter”. For all transcript files, TransDecoder was run using the Universal Genetic Code with a minimum protein length of 100 amino acids “-m 100” in stranded mode “-S”.

Reference protein sequences were obtained from NCBI (O’Leary et al., 2016) from the RefSeq genomes for pea aphids and twenty other invertebrate and vertebrate species [*Aedes aegypti* (GCF_002204515.2), *Aphis gossypii* (GCF_020184175.1), *Apis mellifera* (GCF_003254395.2), *Acyrthosiphon pisum* (GCF_005508785.2), *Bombyx mori* (GCF_030269925.1), *Caenorhabditis elegans* (GCF_000002985.6), *Drosophila melanogaster* (GCF_000001215.4), *Daphnia pulex* (GCF_021134715.1), *Daktulosphaira vitifoliae* (GCF_025091365.1), *Homarus americanus* (GCF_018991925.1), *Homo sapiens* (GCF_000001405.40), *Homalodisca vitripennis* (GCF_021130785.1), *Mus musculus* (GCF_000001635.27), *Macrobrachium nipponense* (GCF_015104395.2), *Myzus persicae* (GCF_001856785.1), *Nilaparvata lugens* (GCF_014356525.2), *Neocloeon triangulifer* (GCF_031216515.1), *Rhopalosiphum maidis* (GCF_003676215.2), *Rhopalosiphum padi* (GCF_020882245.1), *Schistocerca gregaria* (GCF_023897955.1), and *Tribolium castaneum* (GCF_031307605.1)].

Protein files from gene annotation methods of the three assemblies of interest, Trinity-DN, and the RefSeq genomes above were used as input proteomes for ortholog clustering using OrthoFinder v2.5.5 (Emms and Kelly, 2019). OrthoFinder clustered these proteins into so-called orthogroups, which are groups of orthologs inferred to be descended from a single gene in the last common ancestor of all species present in the analysis. This clustering was output in the OrthoGroups.tsv file (see Dryad dataset).

BUSCO v5.2.2 (Manni et al., 2021; Tegenfeldt et al., 2025) was run on the TransDecoder Longest ORFs protein output for PASA, Trinity-GG, StringTie, and Braker for each assembly, as well as Trinity-DN, and the RefSeq proteins. OrthoDB version 10 protein databases for the Eukaryota, Metazoa, Arthropoda, and Insecta levels were used (Tegenfeldt et al., 2025).

### Annotation of genomic interval files

The genomic intervals for mapped transcripts in PASA clusters for each genome assembly were annotated using the OrthoGroups.tsv file output from OrthoFinder, along with header information from the RefSeq protein fasta files for the twenty additional species obtained from NCBI, using custom python scripts. For each potential protein encoded in a transcript in an orthogroup, the first protein listed for a species within the same orthogroup had its header information appended to the corresponding transcript entry in the genomic intervals file as “Species_ortholog:header information”. In cases where a species had no proteins in that orthogroup, the ortholog is listed as ‘none’ in the genomic intervals file. Annotated genomic interval files can be found in the PASA subfolder for each assembly in the Dryad dataset (See **Data Availability Statement**).

### Estimating gene number

The following was performed using custom python scripts. A new orthogroup table was generated replacing the protein(s) (often multiple per locus) from PASA annotations of the three pea aphid assemblies with a single entry for each PASA gene cluster present for each assembly in the orthogroup, along with the proteins present for all other species (and the RefSeq pea aphid proteins). Orthogroups were then analyzed for the number of PASA clusters present for each pea aphid assembly, as well as the number of orthogroup members from the following groups: other pea aphid assemblies (including RefSeq), all non-pea aphid members, as well as all non-aphid members. The total number of PASA clusters for each assembly was then summed based on presence in an orthogroup containing: >1 pea aphid member, ≥1 non-pea aphid member, ≥1 non-aphid member, ≥10 non-pea aphid members, or ≥10 non-aphid members.

### Extracting novel annotations

Orthogroups containing PASA clusters for all three newly annotated assemblies, as well as ≥10 non-aphid members including *Drosophila melanogaster*, but no RefSeq pea aphid member, were extracted from the orthogroup table generated above for further examination using custom python scripts.

### Visualization of results

Plots were generated from comma-separated value files output from custom python scripts visualized in Microsoft Office 365 Excel v2502 and edited in Adobe Photoshop v24.5.0. Gene diagrams were first visualized in Integrative Genomics Viewer v2.16.1 using annotated genomic interval files for PASA clusters in each assembly using a fixed window size across assemblies. Gene annotations were then isolated and compiled in Adobe Photoshop v24.5.0.

## Results

The numbers of transcripts, predicted proteins, proteins in orthogroups, and unassigned proteins, for PASA, Trinity genome-guided, StringTie and Braker for each assembly, as well as for Trinity *de novo* and pea aphid RefSeq annotations, are shown in **Figure 1**. The raw data for this figure can be found in **Supplemental Table S2**. Integration of transcripts from the other four methods into PASA clusters consistently yielded the highest number of transcripts coding for proteins in orthogroups by clustering orthologs of 20 other species with OrthoFinder (**Figure 1 A-C; see Supplemental Table S3 for ortholog clustering**). Braker consistently produced the highest percentage of unassigned proteins of the four genome + RNA-Seq-based methods (1.8%-3.2%) although Trinity *de novo* produced the highest percentage of unassigned proteins overall (5.6%, **Figure 1 A-D, Supplemental Table S2**). Trinity *de novo* also produced the lowest percentage of coding transcripts overall (34.8%), with Trinity genome-guided producing only slightly higher percentages of coding transcripts for each assembly (WD Male Long-Read: 35.8%, WL Male-Long Read: 35.9%, Modified Reference: 36.1%, **Figure 1 A-C, Supplemental Table S2**). Still, both Trinity methods produced a large number of total assembled transcripts relative to other methods, and thus still identified a high number of protein coding transcripts in orthogroups despite the low percentages (**Figure 1 A-D, Supplemental Table S2**). StringTie proved more conservative, consistently predicting much lower numbers of transcripts across each assembly (∼55.9k-58.6k) compared to Trinity genome guided, but with a much higher percentage of coding transcripts (81.6%-87.9%) and the lowest number and percentage of unassigned proteins of any of the methods (144-196, 0.3%-0.4%, **Figure 1 A-C, Supplemental Table S2**).

**Figure 1.**
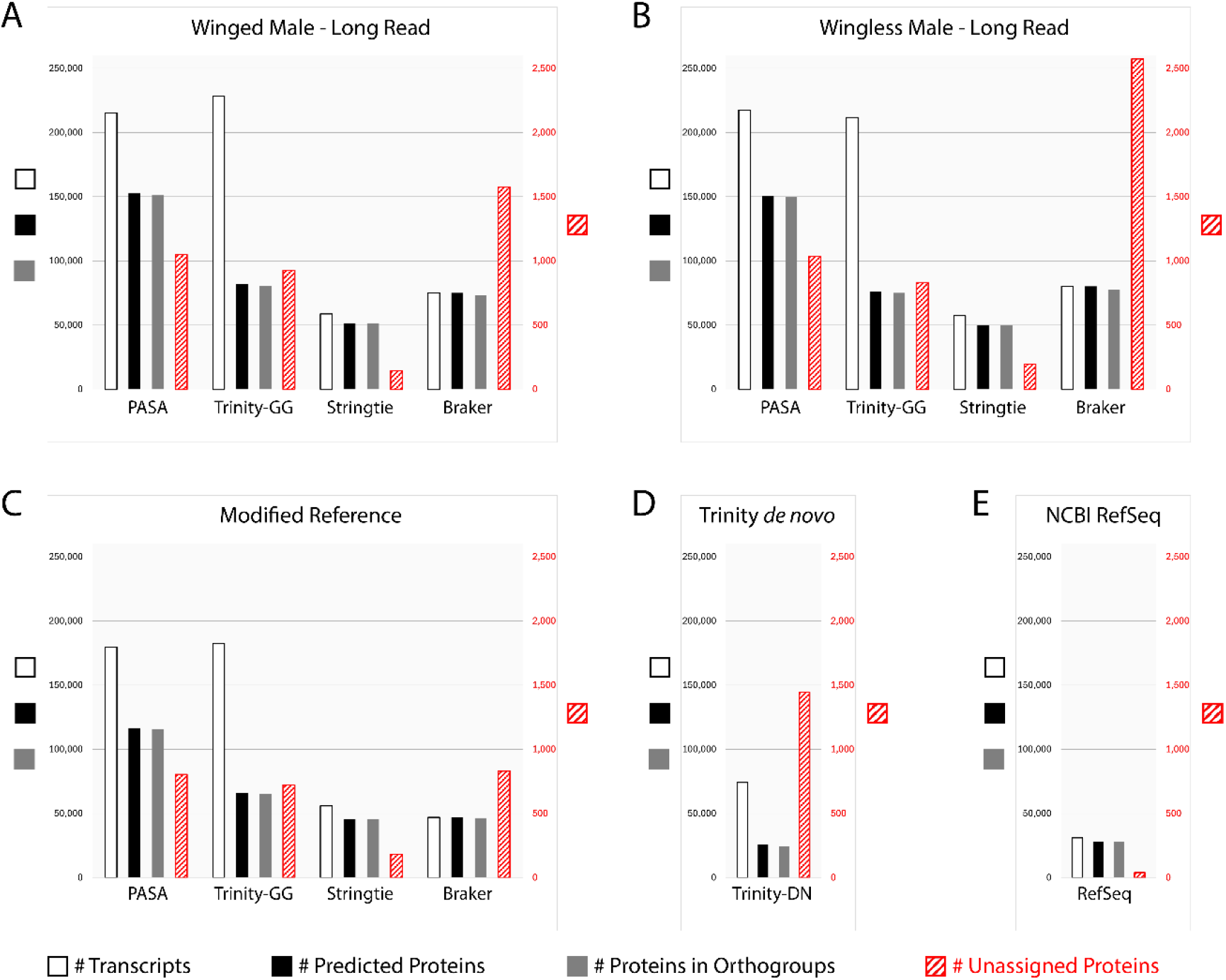
Relative amounts of coding and non-coding transcripts annotated by various methods. Number of predicted transcripts (white), protein-coding transcripts (black), and protein-coding transcripts in orthogroups (gray) are shown on the left axis. Number of predicted transcripts coding for proteins that were not assigned to any orthogroup (red hatched) are indicated on the right axis. All plots share the same scales for both axes. (A-C) Numbers are shown for genome + RNA-Seq-based methods for each assembly. (D) Numbers for Trinity *de novo* assembly of transcripts directly from RNA-Seq reads, integrated with predicted transcripts from the other three methods to produce PASA annotations for each assembly. (E) Numbers for RefSeq annotations of transcripts and proteins in the reference genome.

To assess completeness of the proteomes produced by the various methods outlined above, we employed Benchmarking Universal Single-Copy Orthologs (BUSCO, Manni et al., 2021; Tegenfeldt et al., 2025). We evaluated proteome completeness for all methods, including the RefSeq annotations, at the Eukaryote, Metazoan, Arthropod, and Insect levels using version 10 databases from OrthoDB (Tegenfeldt et al., 2025). The results of this analysis are shown in **Figure 2** (see **Supplemental Table S4** for exact percentages). Integration of transcripts from the four other methods into PASA clusters yielded highly complete predicted proteomes comparable to RefSeq (**Figure 2 A-C, E**). Focusing on the Insecta level, the methods that produced the most complete predicted proteomes were Braker (94.0%-95.6%) and PASA (93.7%-94.9%). The proteomes produced by those two methods were only slightly less complete than the RefSeq annotation (96.7%). This is despite Braker producing the highest percentages of unassigned proteins, likely through detection of Universal Single Copy Orthologs not expressed in our RNA-Seq dataset, which were then incorporated into PASA annotations. StringTie proteomes were less complete than those produced by Braker and PASA (89.6%-91.1%), but still considerably more complete than those produced by Trinity genome-guided (54.0%-55.2%) or Trinity *de novo* (49.7%).

**Figure 2.**
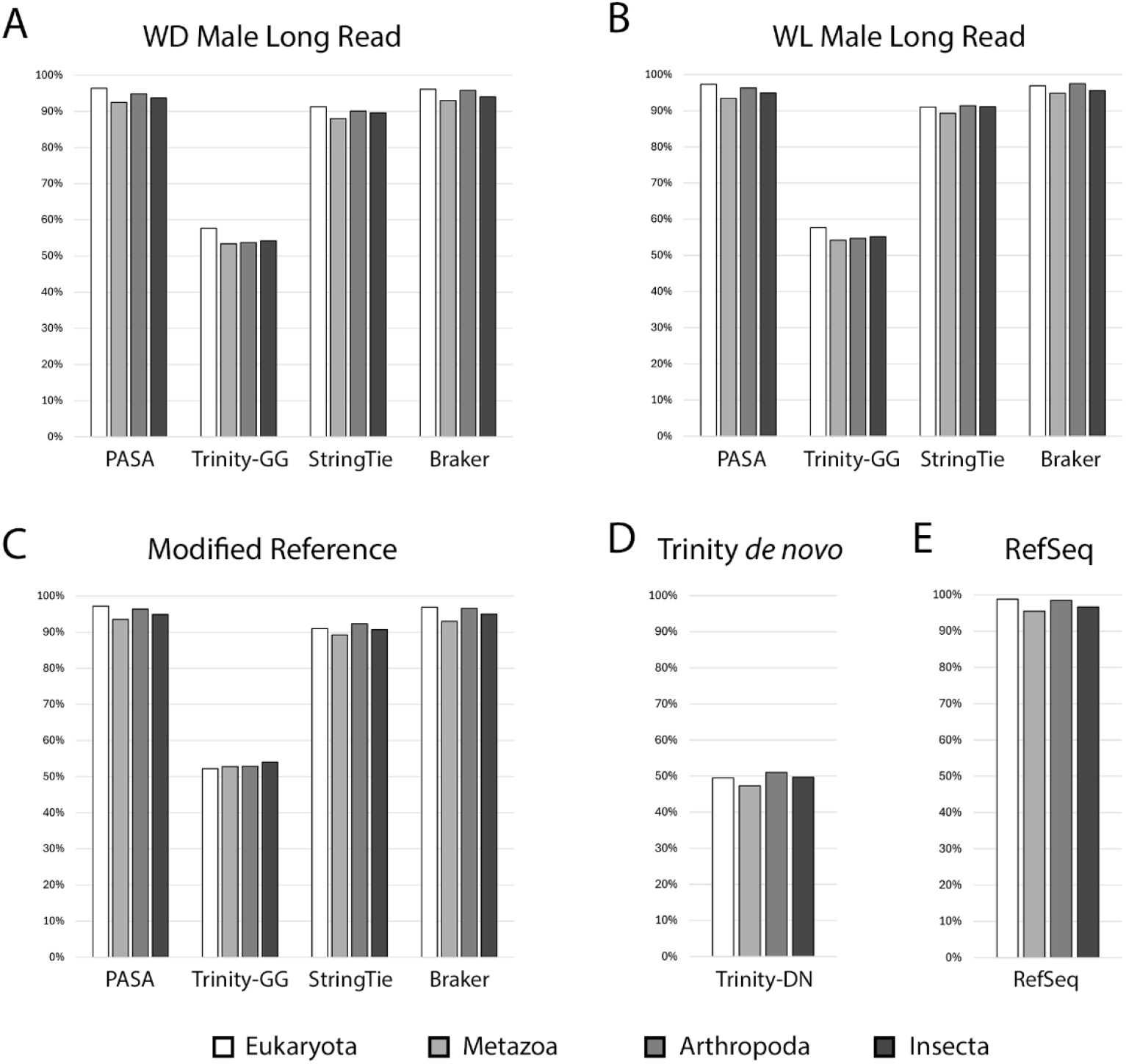
BUSCO analysis of proteomes. The percent of complete BUSCO’s contained in PASA, Trinity genome-guided (Trinity-GG), StringTie, and Braker proteomes for the WD Male Long Read (A), WL Male Long Read (B), and Modified Reference (C) assemblies, as well as Trinity *de novo* (Trinity-DN, D) and RefSeq proteomes (E). Bar shade corresponds to the OrthoDB v10 dataset used: Eukaryota (white), Metazoa (light gray), Arthropoda (gray), or Insecta (dark gray).

Given widespread gene duplication known to occur in pea aphids (Li et al., 2019; The International Aphid Genomics Consortium, 2010), we next asked how many genes are predicted within each of these genomes by our novel annotations. A recent gene number estimate, based on ESTs, cDNA libraries, and WQ-maker predictions, made during assembly of the chromosome-level reference genome, came to 40,090 (Li et al., 2019). We find that our annotation methods produced comparatively high numbers of PASA clusters (141,397 [WD Male LR], 145,815 [WL Male LR], 115,952 [Modified Reference]), likely containing pseudogenes, transposons, and potentially, endosymbiont contamination.

We assessed more stringent gene estimates based on the number of PASA clusters (genes) that code for a protein in an orthogroup with members from other pea aphid assemblies and/or twenty other eukaryotic species (**Figure 3**). Requiring that a PASA cluster from a given assembly codes for a protein in an orthogroup with at least one other protein from another pea aphid assembly produced gene count estimates more similar to the 40,090 estimate of Li et al., 2019 (48,932 [WD Male LR], 47,898 [WL Male LR], 31,888 [Modified Reference]) (**Figure 3, Supplemental Table S5**). The number of genes in orthogroups drops sharply as requirements on the number of orthogroup members from more distantly related species increases (**Figure 3**), suggesting that lineage-specific gene duplicates are rapidly diverging.

**Figure 3.**
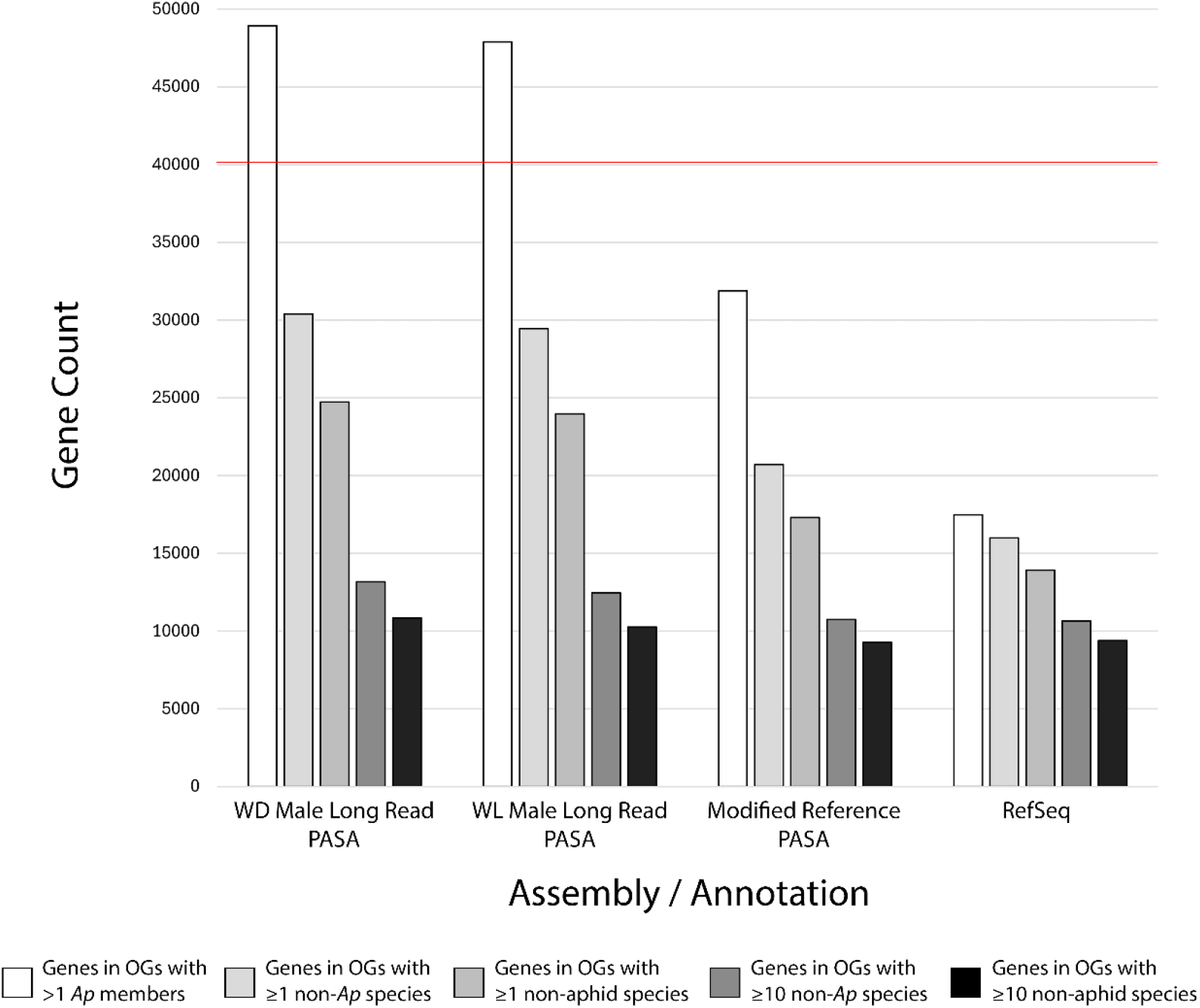
Gene count estimates in pea aphid assemblies with orthologs of varying evolutionary distance. Numbers of PASA gene clusters for each assembly, or RefSeq genes, producing protein-coding transcripts assigned to orthogroups containing members of varying evolutionary distance. The categories are as follows: at least one other member from another pea aphid assembly (white), at least 1 non-pea aphid species (light grey), at least one non-aphid species (medium grey), at least 10 non-pea aphid species (dark gray), or at least 10 non-aphid species (black). The red line indicates the 40,090 gene estimate from the most recent genome assembly (Li et al., 2019).

While a large number of newly annotated genes appear to be lineage-specific, there are also ∼1,000-1,500 more genes with ten or more non-aphid orthologs in the long-read genomes versus the short-read reference (**Figure 3, Supplemental Table S5**). We searched for genes in orthogroups with no RefSeq member but ten or more non-aphid members, including *Drosophila melanogaster*, to identify newly annotated pea aphid orthologs of conserved fly genes. Using this method, we found genes that are missing from the RefSeq annotations (**Figure 4A**), as well as genes whose mis-annotation or mis-assembly disrupted protein clustering (**Figure 4B**). These include missing annotations for *Centrocortin* (*Cen*), *Nad-kinase 2* (*Nadk2*), *Rad17 checkpoint clamp loader component* (*Rad17*), *Cytochrome-c oxidase assembly factor 5* (*COA5*) (**Figure 4A**), and mis-annotation of the genes *trithorax* (*trx*), *Valette* (*Vlet*), and *snail* (*sna*) (**Figure 4B**).

**Figure 4.**
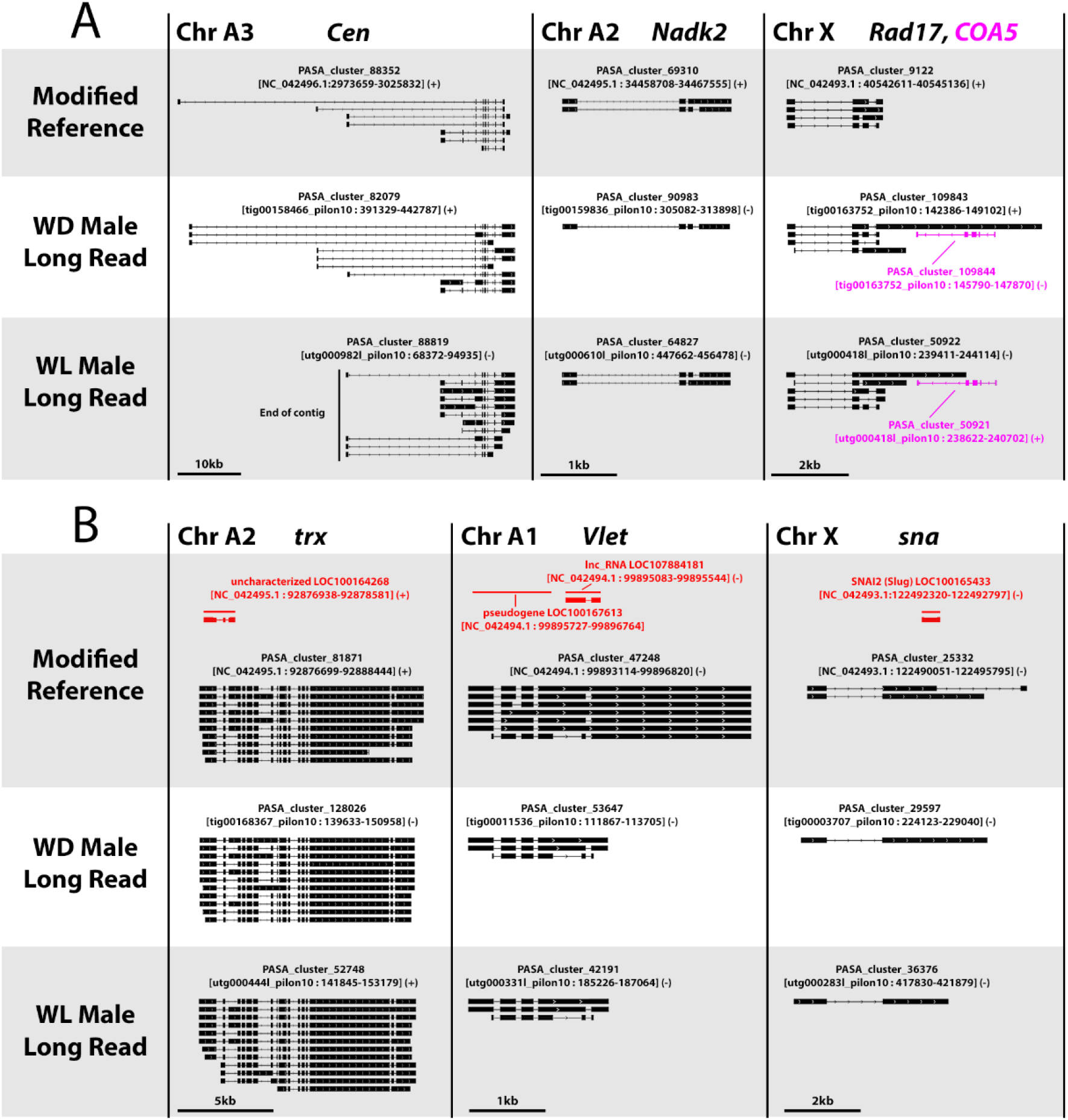
PASA assemblies of missing and mis-annotated genes. Gene structure diagrams for PASA-assembled genes that are either missing or incorrectly annotated in the RefSeq features. RefSeq annotations are shown in red, while newly annotated PASA clusters in the Modified Reference, WD Male Long Read, and WL Male Long Read assemblies are shown in black or magenta. The chromosomal location within the Modified Reference genome is indicated next to the gene name(s). Genes are shown at different scales, indicated by scale bars at the bottom. For clarity, overlapping annotations of other genes are not shown. (A) PASA-assembled gene structures for the genes *Centrocortin* (*Cen*), *Nad-kinase 2* (*Nadk2*), *Rad17 checkpoint clamp loader component* (*Rad17*) (all black), and *Cytochrome-c oxidase assembly factor 5* (*COA5*, magenta) that are absent from the RefSeq annotation. The edge of the contig adjacent to the WL Male Long Read annotation of *Cen* is marked by a black line. (B) PASA assemblies for the genes *trithorax* (*trx*), *Valette* (*Vlet*) and *snail* (*sna*) that are incorrectly annotated in the RefSeq features.

Based on our ability to detect and correct previously unannotated and mis-annotated genes, we checked if these data support the previous identification of four paralogs of the BMP-branch TGF-β ligand *dpp* (Brisson et al., 2010; Saleh Ziabari et al., 2025a). We were able to confirm the presence and gene structure of all four paralogs across all three assemblies (**Figure 5A**). Predicted gene structures, however, did differ between RefSeq features (red) and PASA clusters (black). Our PASA clusters contain novel isoforms using alternative promoters for *dpp-1* and *dpp-3* and predict that the promoter for *dpp-4* is upstream of the RefSeq annotations (**Figure 5A**). These refined promoter annotations for the *dpp* paralogs are consistent across all three genome assemblies.

**Figure 5.**
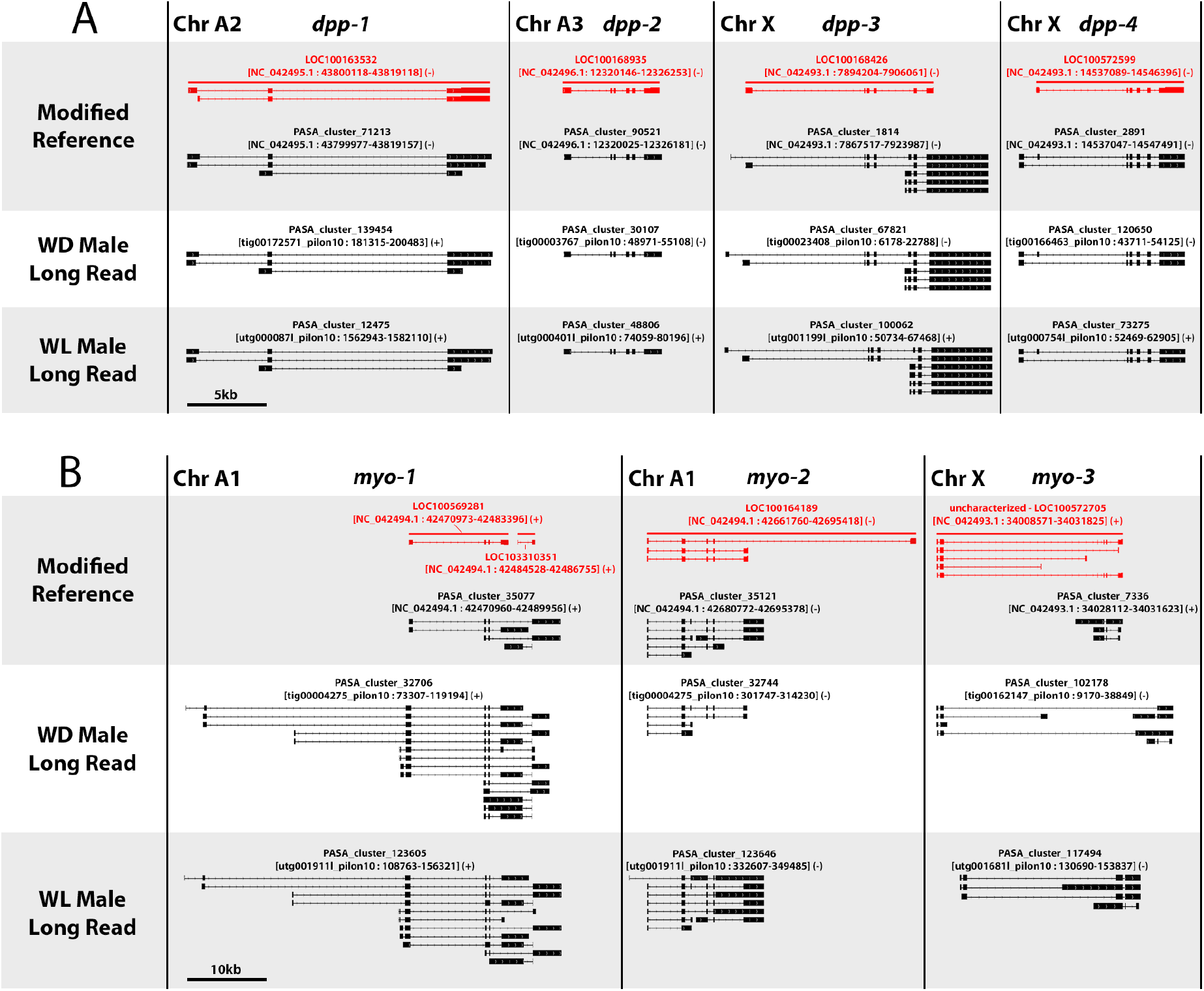
Annotations of duplicated TGF-β signaling ligands. Gene structure diagrams for paralogs of the TGF-β signaling ligands *decapentaplegic* (*dpp*) (A) and *myoglianin* (*myo*) (B) across pea aphid genome assemblies. RefSeq annotations are shown in red, PASA annotations are shown in black. For clarity, overlapping annotations of other genes are not shown. A) Annotations for all four *dpp* paralogs, shown at the same scale. Scale bar: 5kb B) Annotations for the three paralogs of the gene *myo*, shown at the same scale. Scale bar: 10kb

Recent findings have shown that paralogs of the gene *follistatin* are important for pea aphid wing dimorphisms (Li et al., 2020; Saleh Ziabari et al., 2025b). While we have speculated that morph-specific transcripts from *dpp* duplicates may play a role in these dimorphisms (Saleh Ziabari et al., 2025a), potentially interacting with *follistatin* duplicates, Dpp is not known to interact with Follistatin *in vivo* (Bickel et al., 2008; Pentek et al., 2009). For this reason, we set out to identify potential gene duplication for the Activin branch TGF-β ligand and primary Follistatin target, *myoglianin* (Bickel et al., 2008; Pentek et al., 2009). We identified three paralogs of *myoglianin* present in all three assemblies (**Figure 5B**). Two fragmented RefSeq annotations for *myo-1* are actually a single gene with a large duplicated exon containing much of the coding sequence for the separate protein isoforms. PASA clusters in the Long-Read assemblies agree on additional upstream promoters for *myo-1* that are absent from the RefSeq features and PASA clusters in the Modified Reference, indicating potential mis-assembly of this region in the refence (**Figure 5B**). The RefSeq annotation of *myo-2* indicates an alternative downstream exon not supported by RNAseq-based annotation in any of the three genomes (**Figure 5B**). *myo-3* is a pseudogene, with different intact ORFs present in each of the three genome assemblies, responsible for differences in ortholog clustering between them (**Figure 5B**). That is, PASA clusters from the Modified Reference and WD male Long-Read assemblies code for proteins that cluster with other *myoglianin* paralogs and orthologs, but the RefSeq annotation and the PASA cluster in the WL Male Long-Read assembly do not.

## Discussion

Here we present new gene annotations leveraging RNA-Seq data from across life stages, morphs, and sex, for three pea aphid genome assemblies. We integrated annotations from multiple methods (Trinity *de novo*, Trinity genome-guided, StringTie, Braker-ET) into PASA gene clusters, maximizing the number of proteins in orthogroups (**Figure 1**) and overall completeness as determined by BUSCO (**Figure 2**).

Gene count estimates from our annotations of the long-read (48,932 / 47,898) and short-read (31,888) genomes lie above and below, respectively, the previous estimate of 40,090 for the reference genome (**Figure 3, Supplemental Table S5**) (Li et al., 2019). For the long-read genomes, >20k of these genes are predicted to be lineage-specific, while only half as many lineage-specific genes are annotated in the short-read reference (**Figure 3**). This large disparity in gene count is not observed for more broadly conserved genes. New annotations on all three assemblies and RefSeq agree on 9,283-10,854 genes which cluster with orthologs in at least ten non-aphid species (**Figure 3, Supplemental Table S5**). There are likely a number of factors contributing to this large disparity in count estimates for new genes.

The difference in pea aphid gene counts for the reference genome (31,888 in this study, 40,090 in Li et al., 2019) may stem from the numerous differences in estimation methods, as well as our requirement that proteins cluster with those from another pea aphid assembly or the RefSeq proteins. One likely major contributor is that most of the 40,090 predicted genes from Li et al. (2019) do not have associated RefSeq annotations or reference proteins. Additionally, for rapidly evolving gene duplicates that are not part of the reference protein set, mis-assembly in the short-read reference genome may lead to predicted protein sequences that do not cluster with proteins from the properly assembled loci in the long-read genomes. Thus, these mis-assembled genes would not show up in our gene count estimate. Further, even when mis-assembly of duplicates into a single locus in the reference does not disrupt clustering with properly assembled paralogs in the long-read genomes, the gene count estimate for the reference only increases by one, while it increases by more than one for the long-read assemblies.

The two long-read genomes (WD and WL Male) were derived from closely related, inbred lines. Thus, when paralogs are properly assembled in one long-read genome, it is more likely that they will be similarly assembled, annotated and clustered, keeping gene estimates similar. However, there is also the possibility that the gene counts in the long-read genomes (48,932 / 47,898) are artificially inflated by redundant contig fragments. Given these factors, we find it reasonable that we obtained gene counts above and below the 40,090 predicted by Li et al. (2019) for our long-read and reference genomes, respectively. Still, further analyses are required to determine where the true gene count lies within the range shown in **Figure 3**.

The new annotations provided by integrating multiple methods into PASA gene clusters have identified previously missing or mis-annotated genes in the reference assembly (**Figure 4**). These annotations have also helped in confirming previous reports on *dpp* gene duplication (**Figure 5A**) as well as identified a previously unknown *myo* duplicate pseudogene (**Figure 5B**). The new annotations also identify novel promoters for many of these gene duplicates, and consistency of the gene models across the three assemblies strongly suggests that they are accurate (**Figure 5**).

Differences were observed between the assemblies in the annotated gene structures and protein clustering for the *myo3* pseudogene (**Figure 5B**). This seems to be due to differential fragmentation of the ancestral ORF by different premature stop codons in each assembly, disrupting transcript discovery, gene structure annotation, and protein clustering. This highlights an additional challenge in initial pseudogene annotation and detection with a genome assembly generated from a single genetic line. However, such disagreement in structure and coding potential across assemblies may be a hallmark of pseudogenes. Detection of such discrepancies requires a comparative approach, which the data presented here help to facilitate for pea aphids.

These data provide a useful supplement to existing RefSeq annotations by identifying and correcting missing and mis-annotated genes. The annotations of the two long-read genomes provide a platform for investigating mis-assembly of multiple loci into a single locus in the short-read reference genome. These annotations will aid in future investigations into the structure, function, and evolution of genes and gene families.

## Data availability statement

The resources produced in this work have been made available through deposition to public repositories as well as inclusion in the Supplementary data files. The genome assemblies annotated in this work have been deposited to NCBI under BioProjects PRJNA1378164 (Modified Reference), PRJNA1378146 (Wingless Male Long-read), and PRJNA1378148 (Winged Male Long-Read). The male embryo RNA-Seq reads generated in this study were deposited under BioProject PRJNA1380473. Additionally, the predicted transcripts, proteins, and genomic coordinates produced by all methods for all three assemblies, annotated genomic coordinate files for PASA genes, genome sequences with original chromosome names, and the ortholog clustering information for all predicted proteins and those from 20 other species have been deposited to Dryad (https://doi.org/10.5061/dryad.s1rn8pknd).

## Acknowledgements

This research is supported by the National Institute of General Medical Sciences within the National Institutes of Health under award number R35GM144001 to JAB and by the National Science Foundation, Division of Biological Infrastructure, Postdoctoral Research Fellowships in Biology Program under award number 2305817 to KDD.

## Notes

https://doi.org/10.5061/dryad.s1rn8pknd

### Competing Interest Statement

The authors have declared no competing interest.

### Summary of Updates

This version has been revised to update information regarding public submission of the data to NCBI and Dryad, as well as information regarding generation of male embryonic transcriptomes

