## Supplemental Figures and Tables for "New annotations for three pea aphid genome assemblies allow comparative analyses of duplication and gene family evolution": Supplemental_Figures.pdf

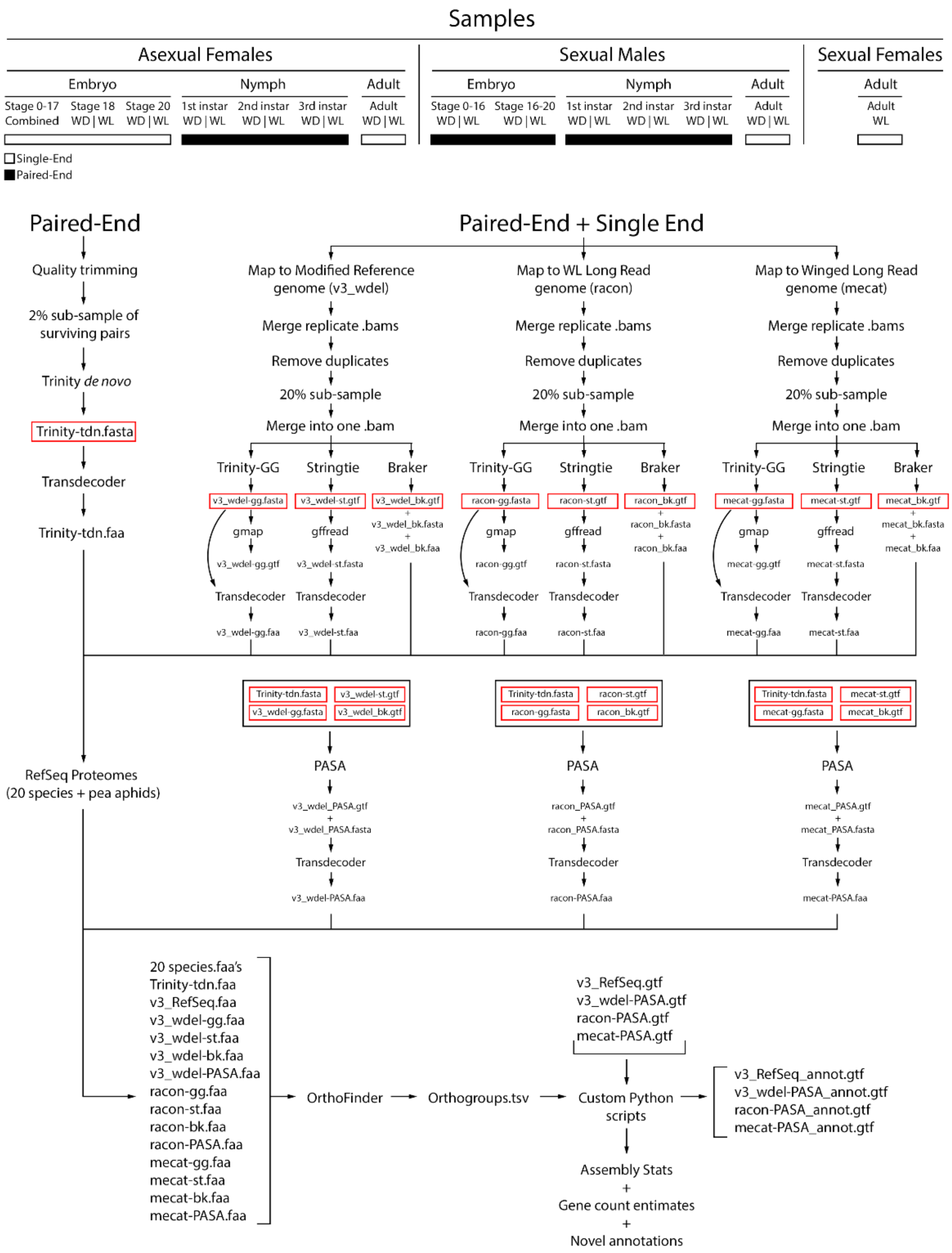


**Supplemental Figure S1. Workflow diagram of genome annotation.**

Information regarding RNAseq samples used in this study are shown at the top with Single-End read samples indicated by a white bar and Paired-End read samples indicated by a black bar. Transcript files from four methods for each assembly used as input to PASA are boxed in red.
